# Glycation and Serum Albumin Infiltration Contribute to the Structural Degeneration of Bioprosthetic Heart Valves

**DOI:** 10.1101/2020.02.14.948075

**Authors:** Antonio Frasca, Yingfei Xue, Alexander P. Kossar, Samuel Keeney, Christopher Rock, Andrey Zakharchenko, Matthew Streeter, Robert C. Gorman, Juan B. Grau, Isaac George, Joseph E. Bavaria, Abba Krieger, David A. Spiegel, Robert J. Levy, Giovanni Ferrari

**Author notes:** Address for correspondence: Giovanni Ferrari PhD - Associate Professor Columbia University, Departments of Surgery and Biomedical Engineering. 630W 168^th^ Street 15.401c New York, NY 10032, T: E.

## Abstract

**Background:** Bioprosthetic heart valves (BHV) are widely used to treat heart valve disease but are fundamentally limited by structural valve degeneration (SVD). Non-calcific mechanisms of SVD entirely account for approximately 30% of SVD cases and contribute to calcific SVD but remain understudied. Glycation mechanisms have not been previously associated with SVD, despite being established as degenerative in collagenous native tissues.

**Objectives:** To determine whether blood component infiltration-based glycation and concomitant human serum albumin (HSA) deposition contribute mechanistically to SVD.

**Methods:** Immunohistochemistry (IHC) was used to identify advanced glycation end-products (AGEs) and serum albumin accumulation in 45 aortic valve BHV explanted due to SVD, glutaraldehyde-treated bovine pericardium (BP) incubated *in vitro* in glyoxal and HSA, and rat subcutaneous BP implants. Structural impacts of glycation-related mechanisms were evaluated by second harmonic generation (SHG) collagen imaging. Hydrodynamic effects of valve glycation and concomitant HSA exposure were studied with an ISO-5840-compliant pulse duplicator system using surgical grade BHV.

**Results:** All 45 clinical explants and in vitro-incubated BP demonstrated accumulated AGE and HSA compared to un-implanted, un-exposed BHV. SHG revealed instigation of collagen malalignment similar to that in SVD explants by glycation and HSA infiltration. Rat subdermal explants also showed AGE and serum albumin accumulation. Pulse duplication demonstrated significantly reduced orifice area and increased pressure gradient and peak fluid velocity following glyoxal and HSA incubations.

**Conclusions:** Glycation and concomitant HSA infiltration occur in clinical BHV and contribute to structural and functional degeneration of leaflet tissue, thus representing novel, interacting mechanisms of BHV SVD.

## Introduction

Bioprosthetic heart valves (BHV) are widely used to treat heart valve disease. BHV leaflets are fabricated from glutaraldehyde crosslinked heterografts—predominantly bovine pericardium (BP) or porcine aortic valve cusps. BHV are increasingly the heart valve replacement prosthesis of choice over mechanical valves due to their technical versatility, adaptability to transcatheter valvular approaches, and freedom from lifelong anti-coagulation therapy^1–3^. However, BHVs are intrinsically limited by degeneration of leaflet tissue matrix, a process named “Structural Valve Degeneration” (SVD). SVD, whose definition has only recently been standardized^3^, constitutes pathologic modification of valve leaflets ultimately compromising biomechanical function. BHV lifespans are limited by SVD^3,4^ to an average between 10-15 years due to functional failure. This process is driven by cellular, biochemical, and biomechanical mechanisms arising from valve properties and design, patient characteristics, and the interactions between them. Calcification is observed in the majority of SVD cases; however, approximately 30% of SVD cases are not associated with significant calcification^5–7^. Further, recent observation suggests that calcification might be associated with SVD without necessarily being functionally causative^5^. Several non-calcific mechanisms have been proposed to contribute to SVD, including inflammatory oxidation, tissue thickening, and collagen network degeneration^4,8^. however, technologies to address known mechanisms thus far have not significantly improved valve durability, suggesting that additional key mechanisms remain unidentified^9,10^.

Protein glycation refers to a complex constellation of non-enzymatic biochemical reactions involving the adduction of sugar-derived moieties to protein nucleophilic groups. Glycation may involve biologically-significant intermediates, such as Amadori products, and culminates in the formation of biologically irreversible advanced glycation end products (AGEs)^11–13^. Glycation is well-established as contributing to structural and functional degeneration in various native tissues and diseases via two major avenues: 1. Direct modification of extracellular matrix (ECM) proteins via crosslinking and modification of protein steric and stereo-electronic interactions; 2. Modulation of cell phenotypes and instigation of inflammation via glycation product-mediated receptor signaling^14,15^. Glycation is known to elicit degeneration of collagenous native tissues via crosslinking and resultant biomechanical deterioration of collagen networks^16–19^; yet, it has not been considered with respect to the pathophysiology of BHV leaflets, for which collagen type I is the majority component^20,21^. Notably, however, a significant difference in the physiochemical mechanisms of glycation in bioprosthetic versus native tissues may arise from the different cellularity contexts. Native tissue glycation is considered to arise primarily from the metabolism of cells resident in the tissue^22^. While attachment and/or infiltration of inflammatory cells has been associated with *de novo* protein deposition in BHVs^23^, they lack living native cells or a functional surface barrier. Therefore, glycation in this context is more likely to occur primarily via infiltration from the surrounding blood of: 1. Glycation precursors that modify the ECM structure directly; 2. Pre-glycated proteins that deposit on or in BHVs; 3. Non-glycated, circulating proteins that are glycated *in situ* by infiltrated glycation precursors. Non-cellular calcium-binding protein adsorption has been proposed in the context of BHV calcifications based on this reasoning^23^. Therefore, in this study, we hypothesized that glycation and infiltration by human serum albumin (HSA), the most abundant and glycation-susceptible circulating protein^24–26^, synergistically contribute to BHV SVD.

## Materials and Methods

### Patient Population

(**Supplemental Table I**). 45 patients with BHV aortic valve replacements requiring reoperation and BHV explant were studied. Patients with endocarditis were excluded. At the initial operation, 25 (55.6%) patients underwent aortic valve replacement only, 6 (13.3%) underwent aortic valve replacement with aortic root replacement, 2 (4.4%) underwent aortic valve replacement with ascending aorta repair, 2 (4.4%) underwent aortic valve replacement concomitantly with CABG, and 10 (22.2%) underwent aortic valve replacement concomitantly with mitral valve repair. BHV explants had implant durations ranging from 3.5 to 14.8 years, with a mean duration of 8.6 ± 0.4 years. Echocardiography preceding reoperation demonstrated BHV aortic insufficiency in 33 (73.3%) of the patients and BHV aortic stenosis in 37 (82.2%) of patients. Comorbidities included aortic dilatation (26.7%), diabetes (17.8%), coronary artery disease (40%), hyperlipidemia (46.7%), hypertension (73.3%), smoking (33.3%), and bicuspid aortic valve (42.2%). Nearly half of patients (44.4%) had been treated with statins.

### Second harmonic generation (SHG) imaging and image analysis

Individual BP discs or valve samples were mounted on an 8mm Fastwell incubation chamber (Electron Microscopy Sciences, Hatfield, PA, USA) and covered with a 0.17 mm coverslip. Second harmonic generation (SHG) microscopy was performed on a Nikon A1RMP multiphoton confocal microscope (Nikon, Minato, Tokyo, Japan). The SHG signal was generated using 860-nm excitation light from a Ti:sapphire femtosecond laser and detected using Nikon Apo LWD 25x/NA1.1 gel immersion as an objective with a 400-450 nm bandpass filter. All images were acquired at 1024×1024 resolution using NIS Elements software. Collagen alignment analysis was performed using CurveAlign software (https://loci.wisc.edu/software/curvealign) and alignment co-efficient was recorded (n=9 images per group)^27^. Collagen crimp distance was measured using NIH ImageJ software (n>50 crimps measured per group).

### *In vitro* glycation/serum albumin exposure

8mm biopsy punches of BP were extensively rinsed in PBS and incubated in either 1x PBS (Corning, NY, USA), PBS + 5% clinical-grade human serum albumin (from stock 25% HSA, Octapharma via NOVA Biologics, Oceanside, CA, USA), PBS + 50mM glyoxal (from stock 88M glyoxal, Sigma Aldrich, St. Louis, MO, USA), or PBS + 50mM glyoxal + 5% serum albumin for 24h, 2 weeks, or 4 weeks at 37°C with 100 rpm orbital shaking. 1mL solution was used per pericardial samples, with 5 specimen/5mL per condition per well in 6-well plates. Plates were sealed with Parafilm and loosely covered with aluminum foil prior to incubation. Subsequent to incubation, BP disks were rinsed in PBS for 5 minutes followed by 3x 1-hour washes in fresh PBS.

### Radiolabeled glyoxal uptake assay

*In vitro* glycation reaction solutions containing 50mM glyoxal with and without 5% bovine serum albumin were spiked with 2.5uCi of 5mCi/mmol ^14^C-labeled glyoxal (American Radiolabeled Chemicals, St. Louis, MO, USA) and applied to 8mm biopsy-punched discs of BP identically to the non-radioactive *in vitro* glycation protocol above for 1,3, 7, 14, and 28 days. Upon harvest, BP discs were rinsed with 1x phosphate-buffered saline at room temperature until the PBS rinse incubated with 5 discs for 1hr achieved background levels of radioactivity according to scintillation counting. BP discs were then desalted by washing 3x in dH2O for 1hr and lyophilized for 2 days, after which their masses were recorded. Lyophilized, weighed BP discs were dissolved in 500uL Biosol (National Diagnostics, Atlanta, GA, USA) at 50° C, followed by addition of 5mL Bioscint (National Diagnostics), mixing and liquid scintillation counting on a Beckman LS-6500 liquid scintillation counter. Scintillation counts per minute were normalized to BP mass, and converted to nanomoles of glyoxal per milligram of dry BP. Preincubation condition were obtained by 24h of incubation with 5% has followed by standard washing with PBS and incubation in 50mM glyoxal in PBS.

### Rat subcutaneous model

8-10 mm discs of BP were rinsed 3 times in 0.9% saline (Rocky Mountain Biologicals, Missoula, MT, USA) and then incubated for 24 hours at 37°C in 0.9% saline. BPs were subsequently rinsed 3 times and stored in 0.9% saline at 4°C until implantation. 3 week-old Sprague-Dawley rats (n=26; Charles River Laboratories, Wilmington, MA, USA) were utilized for all implantation procedures, with strict adherence to an Institutional Animal Care and Use Committee-approved protocol [AAAR6796 (CUIMC)]. All animals were maintained on a standard ad lib laboratory diet. Inhaled isoflurane (Henry Schein Inc., Melville, NY, USA) was used for anesthesia induction (3%) and maintenance (1.5%). Once anesthetized, all animals were weighed and given identifying ear tags. Subcutaneous bupivacaine (6 mg/kg; Auromedics, East Windsor, NY, USA) and meloxicam (2 mg/kg; Henry Schein Inc.) were administered pre-operatively for analgesia. The ventral surface of each animal was clipped and prepped with 70% ethanol and chlorhexidine gluconate (Dyna-Hex 4; Xttrium Laboratories Inc., Mt Prospect, IL, USA). 4 or 6 transverse incisions were made at the dorsal surface, and discrete subcutaneous pockets were formed via blunt dissection. Each animal received one implant per subcutaneous pocket, for a total of 4-6 implants per animal. After ensuring hemostasis, each incision was re-approximated using surgical clips, and the animals were awoken from anesthesia. An additional dose of meloxicam was administered after 24 hours. Surgical clips were removed after 10 days. Animals were sacrificed at either 7 (n=13) or 30 (n=13) days, at which point the implants were harvested and subsequently rinsed and stored in 0.9% saline at 4°C followed by paraffin embedding and sectioning. Blood was drawn at the time of animal sacrifice via cardiac puncture and collected in EDTA blood tubes, after which is was immediately centrifuged at 4°C and 1000 RCF for 15 minutes. The plasma was aliquoted and stored at −80 °C.

### Calcium Analysis

Alizarin Red S (Sigma-Aldrich) staining was used for qualitative calcium analysis, per the manufacturer’s protocol. Quantitative calcium analyses of explanted BP discs were conducted using a colorimetric calcium assay (Sigma-Aldrich), per the manufacturer’s protocol. Explanted BP discs was rinsed in PBS three times and fixed in 10% formalin. Calcium contents were extracted by immersing the fixed discs in 6 M HCl solution overnight at 80°C. The solution was dried overnight at 100 °C and reconstituted in 0.6 M HCl for the colorimetric assay.

### Pulse Duplicator testing conditions

Hydrodynamic pulsatile functionality testing of bioprosthetic valves was performed on a commercial heart valve pulse duplicator (HDTi-6000, BDC Laboratories, Wheat Ridge, CO, USA), which reproduced physiologically equivalent aortic pressures and flows using a PD-1100 pulsatile pump (BDC Laboratories). By measuring the pressure gradient, defined as the difference between the pressures proximal and distal to the valve, the pulse duplicator assesses forward flow through the valve. The pulse duplicator is also used to assess regurgitation, which is an indication of how efficient the valve is at preventing blood from leaking backwards through the valve. Flow, pulse rate, and driving waveform shapes were controlled through Statys^®^ software (BDC Laboratories). The pressure was adjusted via the Systemic Mean Pressure Control knob. All BHVs were subjected to physiological conditions for the aortic position as per ISO-5840^28^ and FDA Replacement Heart Valve Guidance (heart rate at 70 BPM; temperature at 37°C; flow rate at 5L/min; Systolic and diastolic pressures at 110/80 ± 5 mmHg; mean pressure at 95 ± 5 mmHg; systolic duration at 35 ± 5%). Flow, ventricle pressure, and aortic pressure were measured using a transonic sensor (Transonic Systems, Ithaca, NY, USA) and pressure sensors (BDC Laboratories), respectively^29^. Each test run was reported from a 10-cycle measurement average by using Statys™ software (BDC Laboratories), to determine the effective orifice area (EOA) and mean pressure gradient. Valve motion was recorded using two high speed cameras (200 fps, 1280×1024 resolution) at inflow and outflow positions. Peak ejection velocity (V_peak_) was calculated using the following equation, wherein Fmax represents the maximum flow rate and S_max_ represents the maximum actual valve open area. V_peak_ = F_max_/S_max_.

#### a. Surgical aortic valve replacement (SAVR) methods

Three unimplanted Carpentier-Edwards PERIMOUNT RSR surgical bioprostheses (23mm and 27mm, Edwards Lifesciences, Irvine, CA, USA) were used in this study. Upon recording their baseline hydrodynamic functions, two valves were incubated in PBS (23 mm) or 50 mM glyoxal + 5% human serum albumin (27 mm), respectively, at 37°C with 100 rpm orbital shaking. Their hydrodynamic functions were monitored at days 1,3, 7, 14, 21,28, and 35 during the incubation. Valves were placed inside an acrylic fixture set using custom made rubber gaskets, simulating the aortic root and proximal ascending aorta, and a layer of bovine pericardium to prevent paravalvular leak. The fixture was then positioned into the aortic position in the Pulse Duplicator Test System containing 0.9% saline solution. Valves were subjected to physiological aortic conditions as per ISO-5840 and FDA Replacement Heart Valve Guidance (heart rate at 70 BPM; temperature at 37°C; flow rate at 5 L/min; Systolic and diastolic pressures at 110/80 ± 5 mmHg; mean pressure at 95 ± 5 mmHg; systolic duration at 35 ± 5%). Flow, ventricle pressure, and aortic pressure were measured using a transonic sensor (TS410, Transonic Systems) and pressure sensors (BDC-PT, BDC Laboratories), respectively. Each test run was reported from a 10-cycle measurement average by using Statys™ software, to determine the effective orifice area (EOA) and mean pressure gradient. Valve motion was recorded using two high speed cameras (200 fps, 1280×1024 resolution) at the inflow and outflow positions. Peak ejection velocity (V_max_) was calculated by equation below where F_max_ represents the maximum flow rate and S_max_ represents the maximum actual valve open area. V_max_= F_max_/S_max_. For each test condition, the valve loading fixture was disassembled from the test system after one ten-cycle data acquisition and reloaded for additional data acquisition in order to account for potential valve loading artefacts. This process was repeated for 3 10-cycle data acquisition per time point, and values for the 3 repeats per time point were averaged to obtain a data point.

#### b. Transcatheter aortic valve replacement (TAVR) methods

An in-house fabricated self-expanding TAVR valve prototype was used in this study. Upon recording its baseline hydrodynamic functions, the TAVR valve was incubated in 50 mM glyoxal + 5% human serum albumin at 37°C with 100 rpm orbital shaking. The hydrodynamic functions were monitored at days 1, 3, 7, 14, 21, 28, and 35 during the incubation. The TAVR valve was placed inside a conduit fixture set using custom made latex tubing. The fixture was then positioned into the aortic position in the Pulse Duplicator Test System containing 0.9% saline solution. All test conditions were identical to those previously described for the SAVR valve testing.

## Results

*Clinical explant analyses: Calcification, Collagen fiber malalignment, glycation, and serum albumin infiltration in explanted bioprosthetic heart valves are associated with structural valve degeneration.*

With IRB approval [AAAR6796 (Columbia University) and 809349 (University of Pennsylvania)], aortic valve surgical bioprosthetic explants (SAVR) were retrieved for this study (n=45) from patients ranging in age from 34 to 86 years at the time of reoperation (mean = 65.0 ± 2.0 years) (**Supplemental Table I** and **Online Methods and Material**). A transcatheter aortic valve bioprosthesis (TAVR) was also obtained at reoperation and analyzed.

Calcification results (**Table 2**) indicated a wide range of values, with an average leaflet calcification of 125.59 μg calcium per milligram of leaflet mass and a standard deviation of 106.53μg/mg. Calcification results also indicate a large skew, demonstrating an apparently bimodal distribution with 13 valves exhibiting nearly no calcification (<10 μg/mg), only 6 valves with intermediate calcification (between 10-100μg/mg), and 26 valves with high calcification (>100 μg/mg). Unimplanted BHV were initially characterized with second-harmonic generation (SHG) microscopy, which demonstrated an organized alignment of collagen fiber bundles studied (**Figure 1A** and **1B**). Representative microCT and SHG images of explanted clinical BHVs with various degrees of calcification are shown in **Figure 1C** and **1D**. All explanted leaflets, regardless of calcification, showed significant disruption of collagen alignment by SHG (**Figure 1E**) compared to unimplanted BHVs (**Figure 1B**).

**Figure 1.**
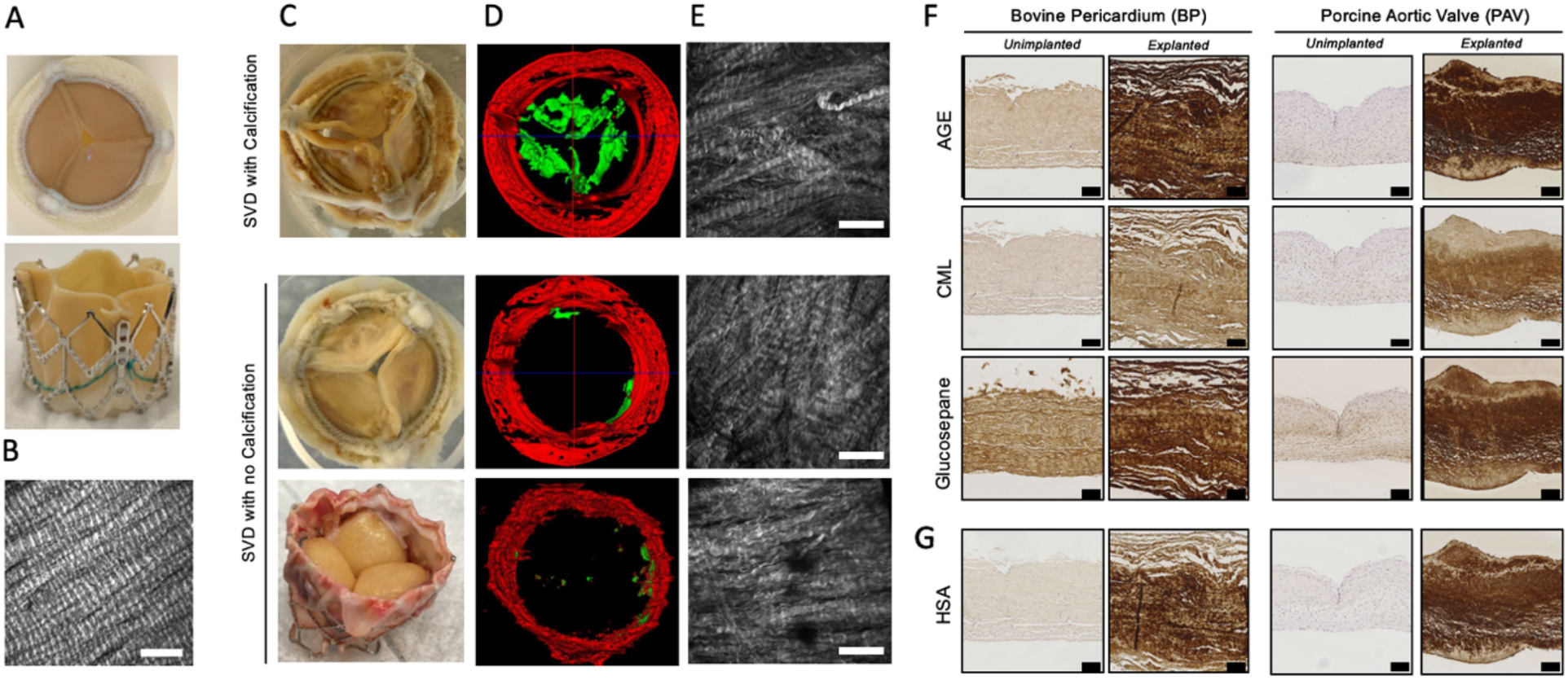
The presence of glycation and serum albumin infiltration in explanted bioprosthetic heart valves (BHV) associated with structural valve degeneration (SVD). **A.** Visual images of unimplanted clinical-grade surgical (Edwards Perimount [Edwards Lifesciences], top panel) and transcatheter (Edwards SAPIEN XT [Edwards Lifesciences], bottom panel) BHV. **B.** Second harmonic generation (SHG) image of collagen fiber bundles in unimplanted BHV leaflet tissue (scale bar represents 100 μm). **C.** Visual images of explanted failed clinical SAVR (top two) and TAVR (bottom). **D.** Micro-computed tomography (microCT) scans, and **E.** SHG images of collagen fiber bundles of calcified (top row) and non-calcified (bottom two rows) valves clinically explanted due to structural SVD (scale bar represents 100 μm). **F.** Immunohistochemistry (IHC) staining for generalized AGE, N-carboxymethyl lysine (CML), and glucosepane in two representatives failed BHV clinical explants alongside unimplanted BHV leaflet tissue for both bovine pericardium and porcine aortic valve BHV (scale bar represents 100μm). **G.** Images of IHC staining for human serum albumin (HSA) from the same valves as **F**.

Clinical explants and unimplanted BHV biomaterials [glutaraldehyde-fixed bovine pericardium (BP) and glutaraldehyde-fixed porcine aortic valve (PAV)] were analyzed by immunohistochemistry (IHC) for generalized AGE, the AGE receptor ligand N-carboxymethyl-lysine (CML)^30,31^, the AGE crosslink glucosepane^32^, and human serum albumin (HSA) (**Figure 1F** and **1G**). Each of the 45 SAVR explants exhibited significant IHC staining for glycation products (**Figure 1F**) and HSA (**Figure 1G**) compared to unimplanted BHVs. IHC mean scores for all 45 explants are shown in **Supplemental Figure 1A.** Statistical analysis revealed no significant relationship of staining intensities to either calcification or the clinical determinants shown, such as diabetes mellitus (**Supplemental Figure 1B**). Collagen malalignment per SHG and positive AGE, CML, HSA, and glucosepane immunostaining were also noted for the TAVR explant (**Supplemental Figure 1C**). In BHV explants fabricated from BP, we observed uniform IHC staining for AGE throughout the tissue, while in BHV explants made from PAV, IHC exhibited nonuniform staining with significant overlap among glycation products and HSA staining patterns (**Fig. 1F-G, Supplemental Figure 1D-E**).

### In vitro model studies of BHV interactions with glyoxal and serum albumin

To investigate the functional mechanisms of glycation and serum protein infiltration, an *in vitro* model utilizing BP, 50mM glyoxal as a glycation precursor^12^, and a physiologic concentration (5% w/v) of clinical-grade serum albumin was designed. IHC on BP following 24 hours of incubation demonstrated glyoxal-generated CML staining and infiltration of HSA uniformly throughout the tissue (**Figure 2A**). Co-incubation with glyoxal and HSA yielded increased CML staining compared to glyoxal alone (**Figure 2A**). ^14^C-glyoxal was utilized in order to measure the glycation capacity of BP. In a 28-day study, uptake approaches an apparent saturation near 30nmol glyoxal per mg tissue, with approximately 50% of the incorporated radioactivity seen at 28 days accumulated within the first 24 hours of exposure (**Figure 2B**). ^14^C-glyoxal incorporation into BP was significantly less in the co-incubated tissue than in BP incubated in glyoxal without HSA (**Figure 2B**). This was expected from inherent competition with the solid BP tissue by dissolved human-serum albumin for reaction with glyoxal; in keeping with this, pre-incubation with HSA followed by incubation in glyoxal alone results in ^14^C-glyoxal incorporation comparable to the glyoxal-only condition. We also sought to visualize whether glycation and HSA infiltration affect the collagen microstructure of BP *in vitro.* In SHG images, BP samples exposed to either glyoxal or HSA for 28 days demonstrated enhanced collagen fiber bundle malalignment and relaxing of crimp compared to BP exposed to phosphate-buffered saline (PBS). This effect was exacerbated in the presence of both glyoxal and HSA (**Figures 2C** and **D**) and comparable with SHG observations of BHV clinical explants (**Figure 1E**). In order to visualize HSA infiltration at the macromolecular level, transmission electron microscopy was performed. Electron micrographs (**Figure 2E**) show longitudinal and cross-sectional views of collagen fibers from individual fiber bundles. BP incubated *in vitro* in 5% HSA with or without glyoxal exhibit an increase in interfibrillar particulates, while incubation with glyoxal alone does not cause any apparent visual difference from PBS after 28 days. In addition, interfibrillar particulates appear closely associated with collagen fiber surfaces and demonstrate non-collagen fibrous aggregates in BP co-incubated with glyoxal.

**Figure 2.**
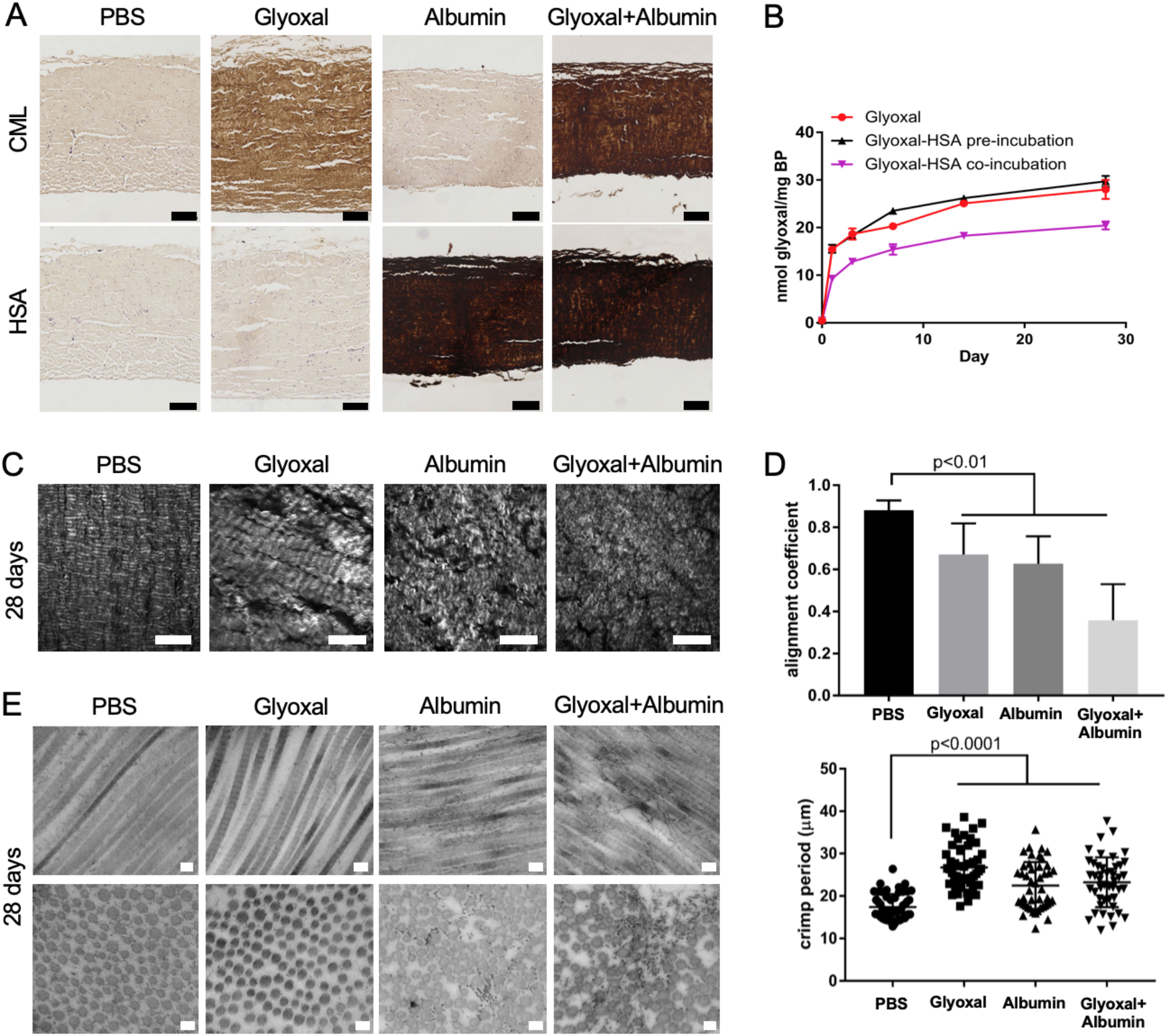
*In Vitro* model studies of BHV interactions with glyoxal and serum albumin. **A.** IHC staining for CML and HSA in cross-sections of glutaraldehyde-fixed bovine pericardium (BP) incubated *in vitro* with glycation precursor and/or HSA for 24 h at 37° C (scale bars represent 100μm). **B.** Time course plot of the uptake of ^14^C-glyoxal by BP over 28 days of incubation in 50mM glyoxal alone with or without 24-hr pre-incubation in 5% HSA or co-incubation in glyoxal and HSA**. C.** SHG imaging of collagen fiber bundles in BP incubated with 50mM glyoxal and/or 5% HSA for 28 days (scale bars represent 100μm). **D.** Quantifications of collagen fiber alignment coefficient (top) and collagen crimp period (bottom) in SHG images of BP after 28-day incubation. **E.** Transmission electron micrographs of longitudinal (top row) and cross-sectional (bottom row) views of collagen fibers in BP incubated for 28 days in PBS or 50mM glyoxal and/or 5% HSA (scale bars represent 100nm).

### Rat subcutaneous explants and circulating biomarkers reveal age-dependent calcification and glycation of glutaraldehyde-fixed bovine pericardium

An established rat subcutaneous implant model (**Supplemental Figure 2A**) of BHV calcification was used to investigate glycation *in vivo* and the impact of animal age in calcification and AGE-mediated SVD^33,34^. Juvenile (3 week-old) and adult (8 month-old) rats received subcutaneous BP implants for either 7 or 30 days^35,36^. Alizarin red staining (**Figure 3A)** revealed diffuse accumulation of calcium phosphate deposits within the subcutaneous explants in juvenile animals, which were more extensive in 30-day explants compared to 7-day explants. No calcification was detectable in adult animals, supporting the clinical observation of age-dependent calcification of BP *in vivo^37–39^.* Quantification of calcium content in BP explants validated this observed increase in calcium accumulation in 30-day explants (159.60 ± 21.17 μg/mg) from 7-day explants (31.88 ± 6.82 μg/mg), both of which demonstrated greater calcium content compared to unimplanted BP (0.36 ± 0.06 μg/mg; p<0.001) and to explants from adult animals (**Figure 3B)**. Established circulating markers of calcification (PO^4-^ [3.004 ± 0.186 vs 2.41 ± 0.120 μmol/mL; p=0.028]; alkaline phosphatase (ALP) [9.192 ± 1.503 vs 0.298 ± 0.130 mU/mL; p=0.004]; and osteopontin (OPN) (33.94 ± 3.54 vs 9.326 ± 0.9261 ng/mL; p<0.0001]) were also elevated in the plasma of juvenile rats when compared to adult animals (**Figure 3C**). SHG analysis of the rat subcutaneous explants **(Figure 3D)** revealed collagen network disruption in cross-sections of both 7- and 30-day BP explants. The alignment coefficients of BP explants were significantly higher in juvenile than in adult animals (0.67 ± 0.02 vs 0.43 ± 0.02; p<0.0001) (**Figure 3E**). In juvenile rats, the 30-day explants demonstrated a higher average crimp distance compared to unimplanted BP (28.68 ± 0.40 vs 25.15 ± 0.45; p<0.0001), indicating the loss of characteristic collagen crimping. Overall, these results indicate progressive and age-dependent glycation and concomitant structural disruption of collagen alignment in BP tissue in the rat subcutaneous implant model. IHC (**Figure 3F**) revealed diffuse CML accumulation within both the 7- and 30-day BP explants that was more prominent in explants from juvenile animals. Plasma concentrations (**Figure 3G**) of sRAGE (3.69 ± 0.465 vs 1.686 ± 0.303 ng/mL; p=0.007), methylglyoxal (3.913 ± 0.875 vs 1.253 ± 0.341 nM; p=0.02), and methylglyoxal protein adducts (6.802 ± 0.758 vs 1.337 ± 0.187 μg/mL; p=0.0001) were all increased in the juvenile rats compared to the adult cohort. IHC using an anti-HSA antibody (**Figure 3H**) that cross-reacts with rat albumin also indicated infiltration of albumin throughout the BP tissue by 7 days, with enhanced accumulation after 30 days. In contrast to all other plasma marker analyses, the plasma concentration of glycated albumin (**Figure 3I**) was lower in the juvenile rats (324.9 ± 45.28 pmol/ml) compared to the adult cohort (539.0 ± 35.37 pmol/ml, p=0.0058). IHC also revealed diffuse staining for OPN (**Supplemental Figure 2B**) and AGE (**Supplemental Figure 2C**) in both 7- and 30-day subcutaneous BP implants, which were qualitatively more pronounced in explants from juvenile animals compared to the adult cohort. IHC for RAGE (**Supplemental Figure 2D**) demonstrated positive staining at the surface of 7- and 30-day explants from both juvenile and adult rats. IHC studies did not detect the presence of glucosepane in the subcutaneous explants (data not shown).

**Figure 3.**
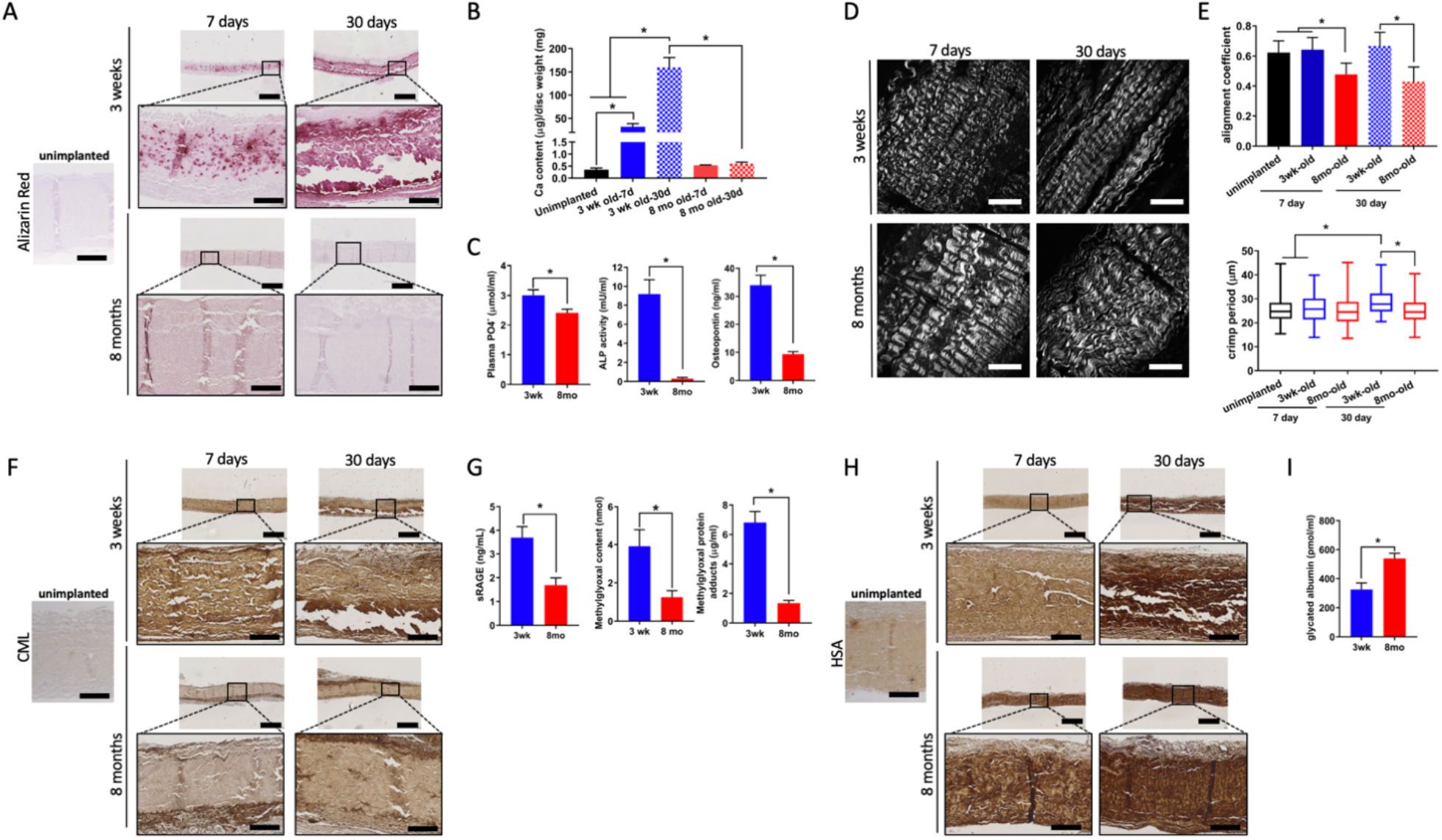
Subcutaneous rat bovine pericardium implants and plasma marker concentrations differ between juvenile and adult models. **A.** Alizarin red staining for calcium on BP following 7- or 30-day subcutaneous rat implantation in 3 week- and 8 month-old rats (scale bars in lower magnification images represent 500 μm; scale bars in higher magnification images represent100μm). **B.** Quantitative calcium analysis from BP explants following 7- and 30-day subcutaneous rat implantation in 3 week-and 8 month-old rats compared to unimplanted BP. **C.** Plasma concentrations of PO_4_^-^, ALP, and OPN in 3 week- and 8 month-old rats. **D.** SHG imaging of collagen fibers in unimplanted and implanted (7- and 30-day subcutaneous implant duration) in 3 week- and 8 month-old rats (scale bars represent 100μm). **E.** Quantification of collagen alignment coefficients and crimp periods in SHG images of unimplanted and implanted BP (7- and 30-day subcutaneous implant duration) in 3 week- and 8 month-old rats. **F.** IHC staining for CML on BP following 7- or 30-day subcutaneous rat implantation in 3 week- and 8-month old rats (scale bars in lower magnification images represent 500 μm; scale bars in higher magnification images represent100μm). **G.** Plasma concentrations of sRAGE, methylglyoxal, and methylglyoxal-derived protein adducts in 3 week- and 8-month old rats. **H.** IHC staining for serum albumin following 7- or 30-day subcutaneous rat implantation in 3 week- and 8 month-old rats (scale bars in lower magnification images represent 500 μm; scale bars in higher magnification images represent 100μm). **I.** Plasma concentration of glycated serum albumin in 3 week- and 8-month old rats.

### Impact of AGE accumulation on bioprosthetic valve hydrodynamic performance

In order to understand the susceptibility of intact BHV leaflets to glycation and human-derived serum albumin infiltration as well as to determine the functional roles of these mechanisms in degeneration of valve performance, we incubated three expired clinical-grade Carpentier-Edwards PERIMOUNT RSR tri-leaflet BP (Edwards Lifesciences, Irvine, CA, USA) BHVs in PBS, 50mM glyoxal in PBS, and 50mM glyoxal plus 5% HSA in PBS, respectively, at 37°C for 35 days and evaluated their hydrodynamic function under physiologic conditions. Hydrodynamic function of the valves was tested at 0, 1,3, 7, 14, 21,28, and 35 days of incubation using an ISO standard heart valve pulse duplicator test system (test conditions in Supplemental Material). The baseline (time point “0”) values of mean pressure gradient and effective orifice area (EOA) of all three valves satisfied the requirements specified in ISO-5840, indicating the expected hydrodynamic performances of SAVR valves used in this study (Figure 4 A-C). Both experimental *in vitro* incubation conditions resulted in a steady decline in EOA (Figure 4 A-C) and increases in pressure gradient and peak jet velocity over time versus PBS incubation. Following 35 days of *in vitro* treatment, the BHV coincubated with glyoxal and HSA demonstrated a 17.48% decrease in EOA, 44.86% increase in pressure gradient, and 7.62% increase in peak jet velocity as compared to each of the baseline values. The BHV treated with glyoxal alone showed a 12.26% decrease in EOA, 27.06% increase in pressure gradient, and 4.95% increase in peak jet velocity as compared to its individual baseline values (Figure 4 A-C). By comparison, the BHV incubated in PBS alone exhibited 4.60%, 15.91%, and 1.99% changes in these three parameters, respectively (Figure 4A to C, Supplemental Figure 3, and Supplemental Table II and III). Energy loss during each cycle was not significantly changed after 35-day PBS incubation (Figure 4D); however, glyoxal and HSA co-incubation resulted in significantly worsened energy loss after 35 days (17.51 ± 0.23 (baseline) vs 30.62 ± 0.35 (day 35), p<0.001) (Figure 4D). SHG imaging (Figure 4E) and analysis (Figure 4F and G) of valve leaflets after 35 days of treatments revealed collagen malalignment and the relaxation of collagen crimp following co-incubated with glyoxal and serum albumin as compared to PBS or glyoxal. Similar to BP glycated *in vitro,* leaflet tissue of the valve treated with glyoxal alone was positively stained for CML, while the valve co-incubated in glyoxal and HSA demonstrated leaflet accumulation of CML and HSA. (Figure 4H and I). We investigated the significance of glycation and concomitant HSA infiltration to TAVR functionality using an in-house fabricated TAVR valve (Supplemental Figure 4A). Valve fabrication is described in the Supplemental Material. Similar to the observations in SAVR valves, TAVR valve also demonstrated collagen malalignment, decline in EOA, and increases in pressure gradient, peak jet velocity, and energy loss (Supplemental Figure 4B-F and Supplemental Table II and III) as a result of co-incubation with glyoxal and HSA. IHC assessments of HSA, AGE, and CML in our fabricated TAVR valve (Supplemental Figure 4C) showed similar results to those in clinical explants as well as our *in vitro* and *in vivo* studies.

**Figure 4.**
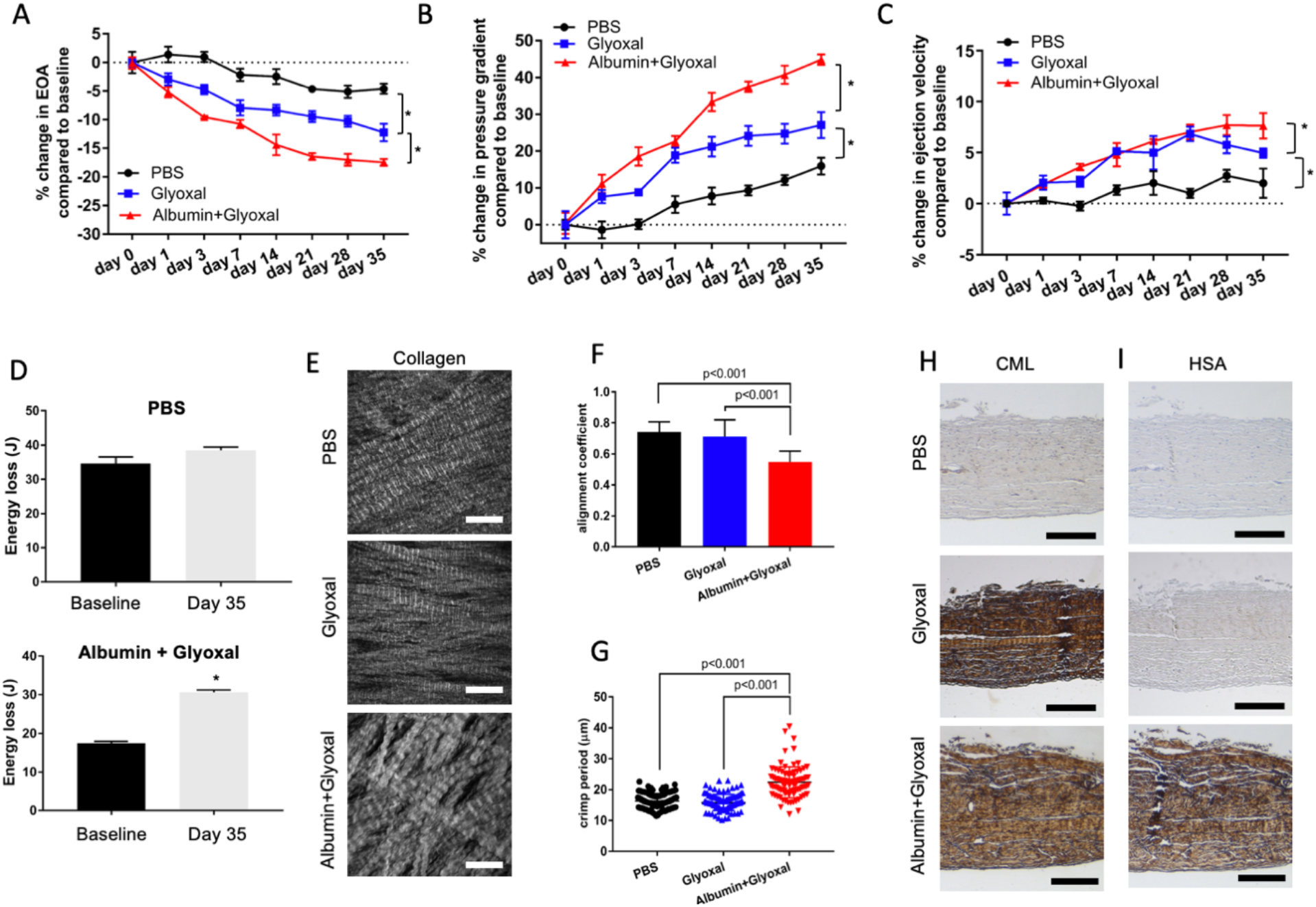
Impact of AGE accumulation on bioprosthetic heart valve hydrodynamic functions. Percent changes in parameters depicting hydrodynamic functions: **A.** EOA; **B.** mean pressure gradient; and **C.** peak ejection velocity, of three surgical aortic bioprosthetic valves (SAVR) measured by a pulse duplicator system during the 35-day *in vitro* incubation period. **D.** Calculated energy loss of BHV before and after 35-day *in vitro* incubation with PBS (top) or HSA and glyoxal (bottom). **E.** SHG imaging of collagen fibers in BHV after 35-day *in vitro* incubation (scale bars represent 100 μm). **F.** Quantification of collagen fiber alignment coefficient from SHG images after 35-day *in vitro* incubation**. G.** Quantification of collagen crimp periods in SHG images after 35-day *in vitro* incubation. **H–I.** IHC staining for CML (**H.**) and HSA(**I.**) in BHV after 35-day *in vitro* incubation (scale bars represent 250 μm).

## Discussion

Glycation is established as a functional mechanism of tissue degeneration in various diseases^8,13,40^. This study is the first description of the involvement of glycation in the degeneration of BHVs. Similarly, while infiltration of circulating proteins on or in clinical^23,41–43^ and *in vivo*^44,45^ implanted BHV tissue has been reported, this work is the first to establish the relevance of glycation and serum albumin infiltration—as well as interactions between them—in BHVs. More importantly, we report that glycation and human-derived serum albumin negatively impact BHV performance. Our study employed a comprehensively translational approach, establishing clinical relevance via explant analyses (**Figure 1**), performing mechanistic modeling *in vitro* (**Figure 2**) as well as *in vivo* (**Figure 3**), and evaluating the functional significance of these mechanisms using a cardiac simulator (**Figure 4**).

IHC on clinical explants indicate that accumulations of AGEs and HSA are prevalent in failed BHVs and show collagen malalignment independently of the extent of calcification. The accumulation of AGEs and HSA, together with the lack of correlation with calcification or any relevant comorbidities tested, suggest that glycation and albumin infiltration are fundamental mechanisms affecting all implanted BHVs.

We sought to understand the mechanisms of AGE and HSA accumulation in BHVs using a glycation/protein infiltration *in vitro* model. The observed accumulation of AGEs and HSA throughout BP tissue after only 24 hours of *in vitro* incubation implies that these intertwined processes begin impinging on clinical BHV essentially immediately upon implantation. Together, clinical correlation and *in vitro* assays suggest that HSA incorporation is enabled by glycation and/or that incorporated HSA increases the tissue’s capacity for glycation. The former possibility is supported by published data suggesting that glycation crosslinking permanently incorporates infiltrated albumin into solid tissue matrices^46–49^. The latter possibility is informed by our *in vitro* modeling observations. While the uptake of radioactive glyoxal seen after serum albumin preincubation is similar to that seen without albumin, these data are normalized to tissue mass without accounting for mass changes due to albumin exposure; as a result, these data may potentially underreport any increase in radioactive glyoxal uptake per unit *tissue* mass due to albumin infusion. Nonetheless, the fact that BP co-incubated with glyoxal and human-derived serum albumin exhibited comparable or greater CML formation after 1-day incubation amid significantly less glyoxal uptake than BP incubated in glyoxal alone carries distinct implications. Glycation occurs primarily on lysine and arginine residues, while the majority of BHV tissue lysines are effectively sequestered from glycation by prior reaction with glutaraldehyde. Incorporation of infiltrated proteins, whose residues are generally unmodified, would increase the repository of glycation-susceptible lysines—as well as arginines—in the tissue. This could explain stronger IHC staining for CML amid diminished glyoxal uptake in albumin co-incubation versus glyoxal alone. These observations suggest that human-derived serum albumin infiltration not only exacerbates accumulation of AGE in BHV, but also modifies the glycation profile toward lysine-directed AGE, which include the most prominent signaling AGE, CML^14,30,31,50^, and the most abundant crosslinking AGE, glucosepane^51–53^. In keeping with this interpretation of mechanistic crosstalk between glycation and serum albumin infiltration, co-incubation led to the highest degree of structural alteration as indicated by SHG assessment and electron microscopy. Additionally, the incorporation of proteins may inculcate BHV tissue with the properties of those proteins, such as calcium- and lipid-binding as well as high oncotic pressure in the case of HSA.

The rat subcutaneous model is an established method for testing BHV biomaterial proprieties and calcification *in vivo.* This study is the first use of this model to investigate glycation in BHV implants and correlate modification of these explants with an assessment of circulating markers. This model resulted in calcium phosphate deposition within the central region of BHV leaflet tissue (comparable to observations in clinical explants^54^), rather than on the surface of the BP tissues, as noted with *in vitro* calcium-phosphate incubations^55,56^. This system also allows modeling major risk factors for BHV failure, such as patient age. The assessments of both explants and circulating markers indicate a host-age dependence of glutaraldehyde-fixed tissue glycation, suggesting that enhanced glycation as well as calcification could contribute to accelerated SVD in pediatric patients. Soluble RAGE (sRAGE) derived from inflammatory cell turnover has been studied as a biomarker for progressive cardiovascular disorders. RAGE is expressed on monocyte cell membranes, and RAGE ligand (such as CML) signaling can initiate monocyte-to-macrophage transition. Inflammatory cell aggregates in BHVs tend to be surface oriented, which is reflected by the hematoxylin counterstain of rat explants. It is possible that AGE formation in BHVs provides the opportunity for RAGE signaling and macrophage deposition to produce reactive oxygen species (ROS) that result in OxAA formation in BHV, as demonstrated by our group in both experimental and clinical BHV studies^5,57^. The lack of demonstrable glucosepane in the rat subcutaneous explants versus virtually all the clinical explants may be due to short (30-day) implantation times relative to the physiologic formation time of glucosepane, which proceeds through long-lived Amadori intermediates^12,53,58^.

Cardiac pulse simulators are required for hydrodynamic performance analysis of prosthetic heart valves, in which pulsatile flow testing is established by the ISO 5840 standard and the U.S. Food and Drug Administration (FDA) guidance. Thus, we aimed to provide functional evidence that AGE and HSA accumulation directly weaken the hydrodynamic performance of BHVs by testing *in vitro*-glycated BHVs. Three key functional parameters, i.e., EOA, pressure gradient, and peak jet velocity, were calculated to assess the hydrodynamic performances of BHV. All three parameters are important indicators for the clinical assessment of aortic valve stenosis severity^28,59–61^. Although the baseline hydrodynamic performances of unimplanted BHV in pulse simulators have been repeatedly reported^62–64^, to the best of our knowledge, we are the first to evaluate the hydrodynamic performances of BHV following *in vitro* incubations that simulate AGE formation observed in clinical explants. In general, all *in vitro* incubation conditions resulted in a steady decline in EOA and increases in pressure gradients and peak jet velocities. Such observations indicate the worsening of the hydrodynamic performances of BHV over time when incubated under simulated physiologic conditions. Glyoxal and glyoxal-HSA co-incubations resulted in pronounced deterioration of hydrodynamic performances as compared to PBS treatment at each time point. These results suggest that the generation of AGEs in BHV significantly alters the biomechanical properties of valve leaflets, potentially causing leaflet stiffening. Additionally, BHV treated with glyoxal alone demonstrated less deterioration of hydrodynamic performances as compared to glyoxal and HSA co-incubation. This could be attributed to the fact that HSA provides additional reactive sites for glyoxal and enhances AGE formation.

### Limitations

There are limitations associated with our study that should be noted. Our clinical series included only aortic valve replacements. Nevertheless, SVD mechanisms, as clinically reported, are comparable regardless of implant site^65,66^. We also restricted our study to a small group of representative glycation-related structures. AGE research has identified a complex myriad of moieties including numerous crosslinks and receptor ligands^11,12^. The HSA used in our *in vivo* studies is a clinical grade, human isolated HSA to closely mimic *in vivo* albumin conditions. Assessing SVD in BP subcutaneous implants that lack exposure to systemic blood flow may be a critical consideration; however, our data show that serum albumin nevertheless permeated throughout the implanted BP as in clinical BHV explants. These studies did not include large animal models involving open heart surgery with BHV valve replacements. These were beyond the scope the present initial discovery investigations. Nevertheless, the rat subcutaneous model has been repeatedly validated is pathologically comparable to circulatory explants in terms of calcification, and the present AGE results from rat explants are comparable to the clinic IHC observations in the present paper. Glucosepane, however, is not detected in subdermal explants, unlike clinical explants; while this could potentially be addressed with longer implantation times, it is a limitation of the current study. There are also some limitations on our pulse duplicator assays including: 1. Using physiologic buffers rather than blood or fluids with viscosity approximating that of blood; 2. The use of clinical BHVs with expired shelf life dates, per FDA requirements, that could alter their susceptibility to glycation. However, the baseline hydrodynamic performances of all these BHVs met the ISO-5840 standards, indicating the satisfactory valve functionality prior to the *in vitro* glycation treatment. It should also be noted that the present studies did not include experiments involving inhibition of glycation, or the use of so-called “AGE-breakers” that have been shown to diminish AGE accumulation experimentally. These studies were not considered because none of the agents previously studied have been shown to be effective in clinical trials, and there are no approved anti-glycation agents available for clinical use. Nevertheless, this first report of AGE and serum albumin accumulation affecting SVD is comprehensive and will provide the basis for therapeutic investigations.

The results of the present studies have important implications for the rapidly increasing use of TAVR. Since its introduction in the early 2000s, TAVR has progressively become an important alternative to open heart valve surgery in patients with high, intermediate, and now low surgical risk and severe aortic stenosis^2–4,67–73^. The recent results of the PARTNER 3^71^ and Medtronic Corevalve Low Risk trial^73^ have significantly altered clinical practice, with TAVR becoming the standard of care for many patients. Yet, it has been simultaneously pointed out that the long-term durability of TAVR due to SVD is currently unclear^74^. In our study, we provided evidence that both clinical grade SAVR and TAVR valves (as well our laboratory-manufactured valves) are similarly subject to glycation and HSA incorporation both *in vivo* and *in vitro.* Furthermore, our pulse duplicator studies show comparable deleterious AGE-related effects on BHVs in flow simulation performance between our TAVR valve prototype and clinical-grade valves. As structural damage associated with leaflet crimping during TAVR valve preparation^75,76^ and increasingly thinner leaflet material^76–78^ are intrinsic characteristics to TAVR versus SAVR valves, it is important to understand the implications of AGE accumulation and microstructural modification on long-term bioprosthesis durability.

Prior research in this field has focused on calcification of BHVs, which has long been associated with the development of SVD. The results of the present studies redirect our current understanding of the pathophysiological mechanisms of SVD by addressing AGE and serum protein mechanisms. Based on these data we propose that glycation and protein infiltration result in BHV tissue matrix disruption via multiple mechanisms: 1. Reduction of BHV leaflet mechanical compliance due to AGE crosslinking; 2. Enabling of the permanent incorporation of infiltrated proteins, and their properties, such as calcium or lipid binding, via AGE crosslinking^46–49^; 3. Modification of BHV leaflets by AGE and protein incorporation that can alter collagen fiber interactions and resultant force dissipation during biomechanical activity^16–18^; 4. Pro-inflammatory responses to the valve leaflet tissue by signaling to receptors of glycation products, including the receptor of AGE (RAGE)^14,15,31^. Glycation-based permanent incorporation of infiltrated proteins is expected to disrupt BHV biomechanical functionality on the macroscopic scale while disrupting tissue matrix structure on the microscopic scale. Thus, it is concluded that the accumulation of AGE and serum albumin in clinical explants and the impact of glycation on both the collagen fiber microstructure and on the hydrodynamic function of BHVs significantly contribute to SVD. (**Figure 5**).

**Figure 5.**
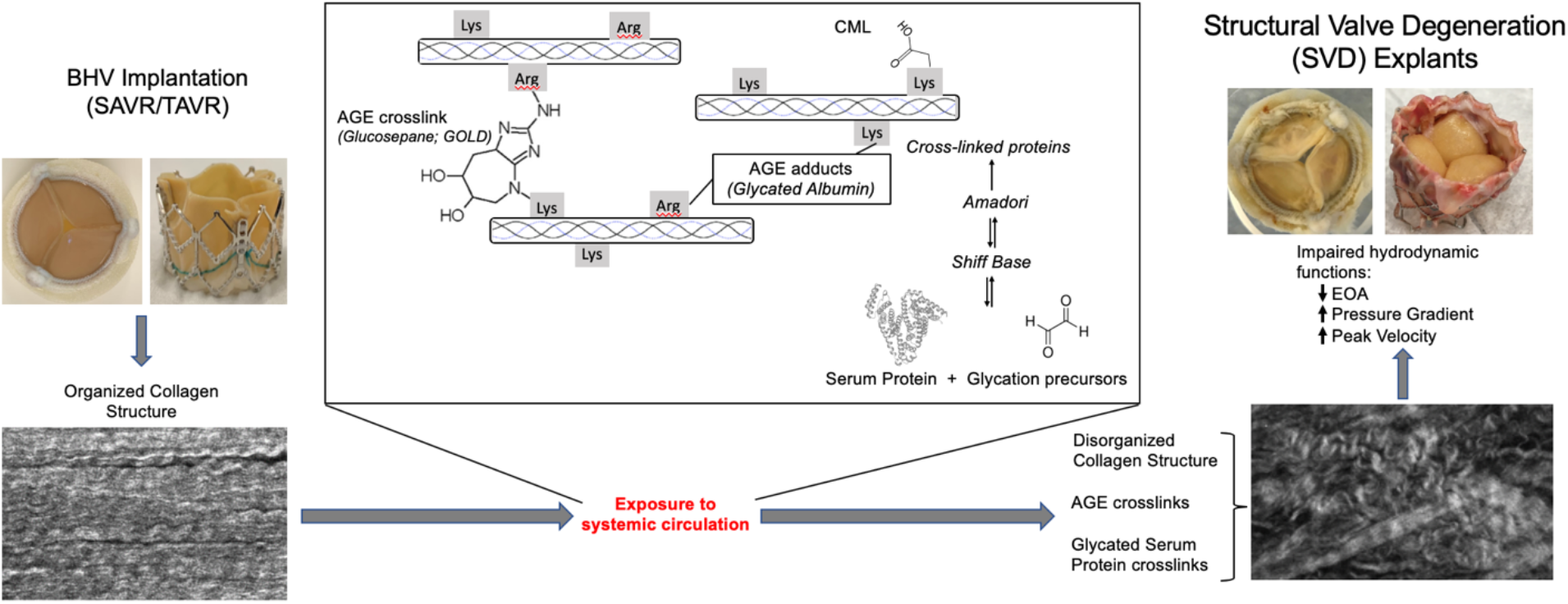
Conceptual model of the synergistic effects of glycation precursor and protein infiltration on BHV structure and function during clinical implantation. BHV exhibit structurally undisrupted, biomechanically functional leaflets and organized collagen microstructure at clinical implantation. Exposure to systemic circulation during implantation causes the progressive infiltration of glycation precursors and serum proteins. Glycation causes biochemical modification of the BHV tissue matrix, including: crosslinking, the generation of inflammatory stimulators, and the permanent incorporation of infiltrated proteins. Tissue protein modification and the incorporation of circulating proteins from the patient drive collagen disorganization and biomechanical functional disruption in the BHV tissue.

## Author contributions

AF, RJL, and GF conceived and planned the experiments. GF and RJG secured funds. AF, YX, APK, SK, CR and AZ carried out the experiments. IG, JEB, MS, and DAS contributed to surgical implants, explants, and non-commercially available supplies and their usage. JBG contributed to the interpretation of the results. AF, RJL and GF wrote the manuscript. All authors provided critical feedback and helped shape the research, analysis and writing the manuscript.

## Supplementary Materials

### Materials and Methods

#### Clinical explant processing

Clinically-explanted surgical bioprosthetic aortic valve replacements were retrieved under IRB approvals [AAAR6796 (Columbia University) and 809349 (University of Pennsylvania)]. Explanted BHV were rinsed in PBS before fixation in 10% neutral-buffered formalin for 24-to-72 hours, followed by dehydration in graded ethanol and storage in absolute ethanol at 4°C. Individual intact leaflets were excised, cut in half vertically from free edge to base, and embedded cut-edge-down in paraffin blocks for sectioning and histological analysis

#### Bovine pericardium processing

Intact, de-fatted, and cleaned pericardia from freshly-slaughtered mature bovines were obtained on ice from Animal Technologies, Inc (Tyler, TX, USA). Pericardial sacks were immersed in a 0.6% glutaraldehyde, 50mM HEPES pH 7.4 solution for 7 days at room temperature in capped 1L glass Corning bottles. Next, pericardia were rinsed in cold saline and stored in 0.2% glutaraldehyde, 50mM HEPES pH 7.4 solution at 4° C in capped 1L glass Corning bottles.

#### Plasma Analysis

Rat plasma phosphate (Abcam; Cambridge, UK), alkaline phosphatase (Abcam), osteopontin (R&D Systems; Minneapolis, MN, US), soluble receptor for AGEs (sRAGE; Abcam), glycated albumin (G-Biosciences; St. Louis, MO, USA), methylglyoxal (Abcam), and methylglyoxal protein adduct (Abcam) concentrations were measured by corresponding commercially-available assay kits, respectively, per manufacturer protocols.

#### Immunohistochemistry (IHC)

Excised leaflets from explanted clinical BHVs and clinical BHVs used for *in vitro* experiments, unimplanted BHV biomaterial controls, rat subcutaneous BP explants, and *in vitro*-glycated BP samples were fixed in 10% formalin for 24-72hrs and embedded in paraffin. 5μm-thick sections were cut via microtome and mounted on high-adherence glass microscope slides. Antigen retrieval was accomplished by overnight incubation in 1x sodium citrate buffer (Sigma Aldrich St. Louis, MO, USA) at 60°C. For glucosepane IHC, antigen-exposed sections were incubated for 24hr in 100mM sodium phosphate buffer pH 9.0 + 80mM NaBH4 in order to reduce artefactual moieties generated by glutaraldehyde fixation before primary antibody incubation. Anti-AGE (Abcam, Cambridge, UK #ab23722, 1:5000 dilution), anti-CML (Abcam #ab27684, 1:5000 dilution [1:1000 dilution for rat subcutaneous explants]), anti-HSA (Abcam #ab117455, 1:100 dilution [1:1000 dilution for rat subcutaneous explants]), anti-osteopontin (Abcam #ab8448, 1:2000 dilution), anti-RAGE (Abcam #ab3611, 1:2000 dilution) and anti-glucosepane (David Spiegel laboratory, Yale University, New Haven, CT, per material transfer agreement, 1:100 dilution) antibodies were diluted in DAKO primary antibody diluent (Agilent Technologies, Santa Clara, CA, USA) and applied to sections overnight (16-18hrs) at 4°C with gentle orbital shaking. DAKO HRP polymer-conjugated anti-rabbit and anti-mouse secondary antibodies (Agilent, #ab214880, #ab214879) were applied at room temperature for 1hr. Washes were accomplished using 1x dilution of DAKO Wash Buffer 10x (Agilent). IHC stains were developed for 8 minutes using 3,3’-diaminobenzidine tetrahydrochloride substrate (Abcam) and sections were counterstained for 3 minutes with Gill’s haematoxylin (Sigma-Aldrich, St. Louis, MO, USA). Slides were mounted using Permount mounting medium (Fisher Scientific, Hampton, NH, USA) and visualized by 20x light microscopy and 4-40x Aperio (Leica Camera AG, Wetzlar, DE) scanning. Intensity of IHC staining was evaluated by 4 independent judges (authors AF, SK, RL, and GF) according to the following scoring system: 0 = Trace or no staining; 1 = Light or highly spatially-restricted staining; 2 = Moderate staining; 3 = Strong staining with good coverage; 4 = Extremely strong and/or complete staining. The averages of the four judges’ scores were taken as the overall scores for each stain for each valve. Statistical comparisons were performed using two-tailed Student’s T-tests and Wilcoxon rank sum tests for ordinal data and Pearson’s and Spearman’s correlations for continuous data (calcification).

#### Fabrication of Percutaneous Bioprosthetic Heart Valve

Each stent is manufactured out of a drawn Nitinol tube of 7mm diameter and 0.50 mm wall thickness. The tube is fed into a laser cutting machine, which cuts out the specified stent pattern accordingly. A de-burring step allows for removal of small parts and chips of Nitinol that were not cut completely during the laser cutting step before the critical shape-setting step, which produces the stent’s expanded size and shape. Nitinol’s super-elastic properties^79^ allow for shaping the cut out tubular stent from a 7mm diameter stent to a 21-23 mm diameter stent without compromising its structural integrity. Once the shape-setting step is completed, the stent is brought into the electropolishing station where the stent is dipped into a variety of baths containing proprietary solutions that electropolish the surface of the stent to optimize its biocompatibility and host tissue response once implanted. Throughout the manufacturing of the stents, various inspection steps are performed to ensure the stent’s final dimensions, structural integrity, and surface finish quality. Three separate pieces of pericardium are cut to specified dimensions and geometries. The three sections are then sutured together with 1 mm of overlap using 5-0 non-absorbable monofilament sutures to form an appropriately size pericardial conduit. The conduit is then attached to the stent with temporary sutures. Using a continuous 5-0 leaflet suture with dual needles the conduit is permanently fixed to the stent by following the leaflet contour around the stent struts. One suture with dual needles is used to ensure a continuous thread throughout the valve contour and eliminates knots which can negatively impact valve performance and durability at these critical areas. With each needle end the leaflets are then fixed to the commissural posts using an in and out suturing technique around each post. A continuous 6-0 suture connects the valve conduit inflow region to the inflow edge of the stent. Once the inflow attachment is completed the excess pericardium is trimmed appropriately.

#### Transmission electron microscopy

BP discs were incubated in PBS (Corning, NY, USA) or PBS + 50mM glyoxal (from stock 88M glyoxal, Sigma Aldrich, St. Louis, MO, USA), and/or 5% HSA (from stock 25% HSA, Octapharma via NOVA Biologics, Oceanside, CA, USA) for 28 days at 37°C with 100 rpm orbital shaking. Post-incubation, standard PBS washing was performed before discs were fixed with 2.5% glutaraldehyde and post fixed in 1% osmium tetroxide for 1 hour, dehydrated with alcohol and infiltrated in epoxy resin overnight, and cured for 48 h in a 60°C oven. The resulting resin blocks were trimmed with a razor blade into a trapezoid block face such that the y-axis represents the transmural/thickness axis and the x-axis represents the circumferential axis. Thin sections (60 nm) were cut using a Diatome Ultra Diamond Knife 35° (Diatome Diamond Knives, Hatfield, PA, USA), picked up on grids, which were then stained with uranyl acetate and lead citrate. Samples were then examined using a JEOL JEM-1200 EXII electron microscope (JEOL USA, Peabody, Massachusetts, USA). Images were captured at 100,000x magnification with an ORCA-HR camera (Hamamatsu Photonics, Bridgewater Township, NJ, USA) and recorded with AMT Image Capture Engine.

---

**Supplement Table 1.** Patient Demographics and Clinical Characteristics of Bioprosthetic Aortic Valve Explants

| **Summary and Demographics** |  |  |  |
| --- | --- | --- | --- |
| **Total, n** | **45** | **Diagnosis, n (%)** |  |
| Male, n (%) | 28 (62.2) |  |  |
| Female, n (%) | 17 (37.8) | AV Insufficiency | 33 (73.3) |
| Age (Range), years | 64.96 (34 - 86) | AV Stenosis | 37 (82.2) |
| **Comorbidities, n (%)** |  | AV Calcification | 21 (46.7) |
|  |  | Dilated Aorta | 12 (26.7) |
|  |  | MV Insufficiency | 33 (73.3) |
| Smoking | 15 (33.3) | Aortic Peak Gradient > 40 mmHg | 36 (80.0) |
| Diabetes Mellitus | 8 (17.8) | **Procedure, n (%)** |  |
| Hypertension | 33 (73.3) |  |  |
| CAD | 18 (40.0) | AVR Only | 25 (55.6) |
| History of CABG | 15 (33.3) | AVR + Aortic Root Replacement | 6 (13.3) |
| Hyperlipidemia | 21 (46.7) | AVR + CABG | 2 (4.4) |
| Peripheral Arterial or Cerebrovascular Disease | 10 (22.2) | AVR + MVR | 8 (17.8) |
| Previously BAV | 19 (42.2) | AVR + MVR + CABG | 2 (4.4) |
| Creatinine > 1 mg/dL | 17 (43.6) | AVR + AA Repair | 2 (4.4) |
| Creatinine > 2 mg/dL | 3 (7.7) | **Bioprosthesis Size (mean duration, years), n (%)** |  |
| Statin Therapy | 20 (44.4) |  |  |
| NYHA Class |  | 19 mm (9.0 ± 0.77) | 4 (8.9) |
| 1 | 3 (6.7) | 21 mm (5.9 ± 0.25) | 9 (20.0) |
| 2 | 16 (35.6) | 23 mm (8.9 ± 0.56) | 12 (26.7) |
| 3 | 18 (40.0) | 25 mm (8.3 ± 0.49) | 9 (20.0) |
| 4 | 8 (17.8) | 27 mm (9.6 ± 0.52) | 5 (11.1) |
| **Bioprosthesis Used, n (%)** |  | 29 mm (10.6 ± 0.37) | 6 (13.3) |
| St. Jude (Porcine) | 7 (15.6) |  |  |
| Medtronic (Porcine) | 2 (4.4) |  |  |
| Carpentier-Edwards (Bovine) | 31 (68.9) |  |  |
| Carpentier-Edwards (Porcine) | 1 (2.2) |  |  |
| Sorin (Bovine) | 4 (8.9) |  |  |

**Supplement Table II.** Statistical analysis of the three tested hydrodynamic parameters (Effective orifice area (EOA), pressure gradient, and ejection velocity) of SAVR and TAVR valves.

|  | Mean 35 days | Mean baseline | Standard deviation | P-value |
| --- | --- | --- | --- | --- |
| SAVR (Glyoxal) | 2.100 | 2.393 | .0294 | .0003 |
| SAVR(Albumin + Glyoxal) | 2.420 | 2.933 | .0222 | <.0001 |
| SAVR (PBS) | 2.227 | 2.327 | .0379 | .0318 |
| TAVR (Albumin + Glyoxal) | 1.520 | 1.727 | .0129 | <.0001 |

**PRESSURE GRADIENT**
|  | Mean 35 days | Mean baseline | Standard deviation | P-value |
| --- | --- | --- | --- | --- |
| SAVR (Glyoxal) | 7.200 | 5.667 | .2040 | .0008 |
| SAVR (Albumin + Glyoxal) | 5.867 | 4.050 | .0856 | <.0001 |
| SAVR (PBS) | 5.100 | 4.333 | .0917 | .0003 |
| TAVR (Albumin + Glyoxal) | 13.000 | 9.933 | .1291 | <.0001 |

**EJECTION VELOCITY**
|  | Mean 35 days | Mean baseline | Standard deviation | P-value |
| --- | --- | --- | --- | --- |
| SAVR (Glyoxal) | 1.609 | 1.533 | .0126 | .0018 |
| SAVR (Albumin + Glyoxal) | 1.333 | 1.239 | .0112 | .0005 |
| SAVR (PBS) | 1.382 | 1.356 | .0142 | .0811 |
| TAVR (Albumin + Glyoxal) | 1.335 | 1.272 | .0119 | .0028 |

**Supplement Table III.** Statistical analysis comparing the effect of *in vitro* treatment on the three hydrodynamic parameters (EOA, pressure gradient, and ejection velocity) of SAVR valves.

|  | Mean Treatment | Mean PBS | Standard deviation | P-value |
| --- | --- | --- | --- | --- |
| Glyoxal EOA | 2.100 | 2.227 | .0294 | .0062 |
| Glyoxal Pressure | 7.200 | 5.100 | .1581 | <.0001 |
| Glyoxal Velocity | 1.609 | 1.382 | .0146 | <.0001 |
| Albumin+ Glyoxal EOA | 2.420 | 2.227 | .0191 | .0002 |
| Albumin+ Glyoxal Pressure | 5.867 | 5.100 | .0817 | .0003 |
| Albumin+ Glyoxal Velocity | 1.333 | 1.382 | .0176 | .0269 |

**Supplemental Figure 1.**
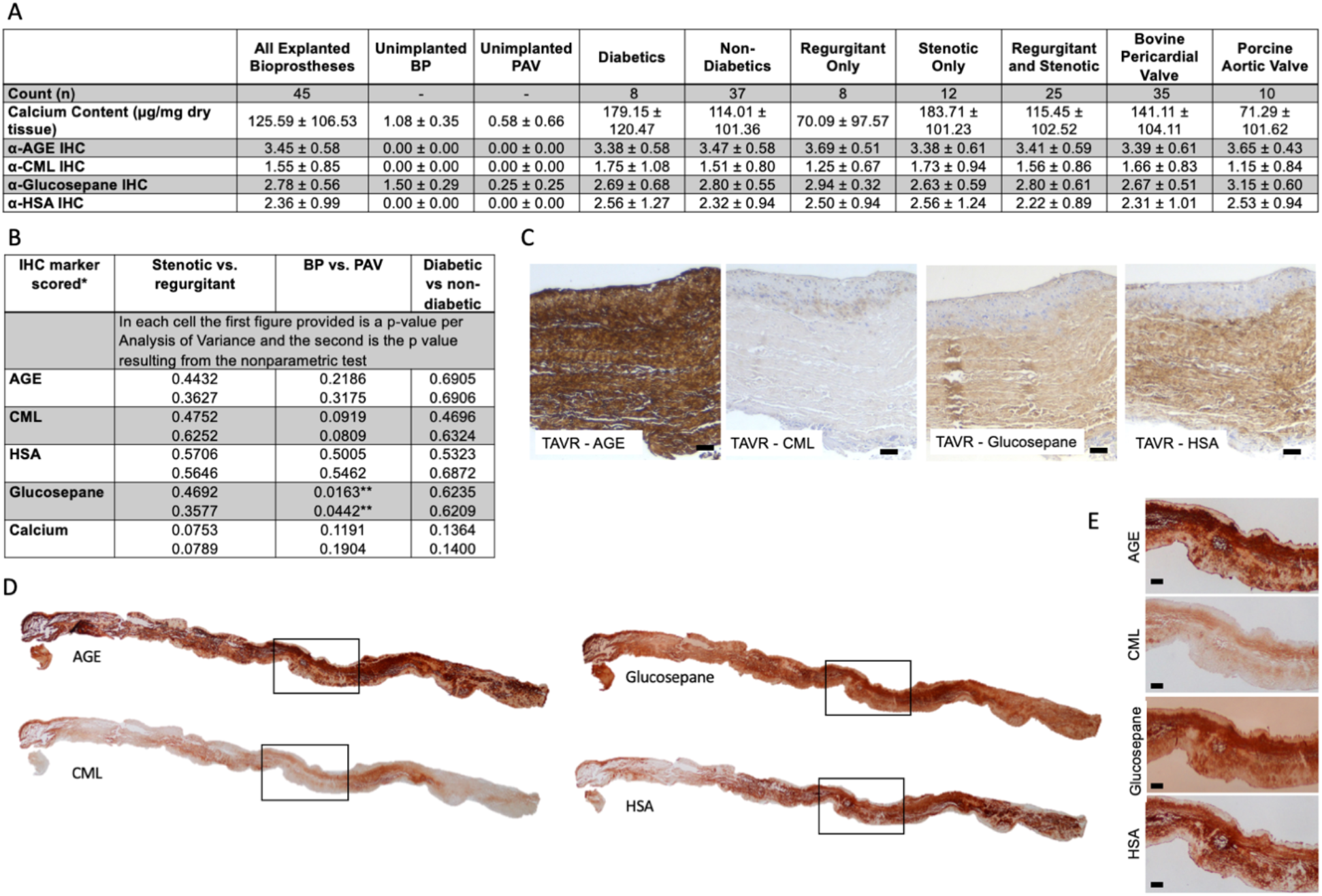
Quantifications, statistics, and additional observations from clinical explant studies. **A.** Quantitative calcium content data and semi-quantitative IHC scoring data for the 45 clinically-explanted BHVs included in the study binned as indicated. **B.** Statistical analysis of select comparisons from the data presented in **A**. **C.** IHC stains for generalized AGE, CML, glucosepane, and HSA in a clinical TAVR valve explanted after 1 year (scale bars 50um). **D.** Compiled micrographs of full-length PAV explant sections IHC stained for indicated targets. Boxes indicate regions expanded in **E**. **E.** Zoomed-in views of section regions indicated by boxes in **D** (scale bars represent 100um).

**Supplemental Figure 2.**
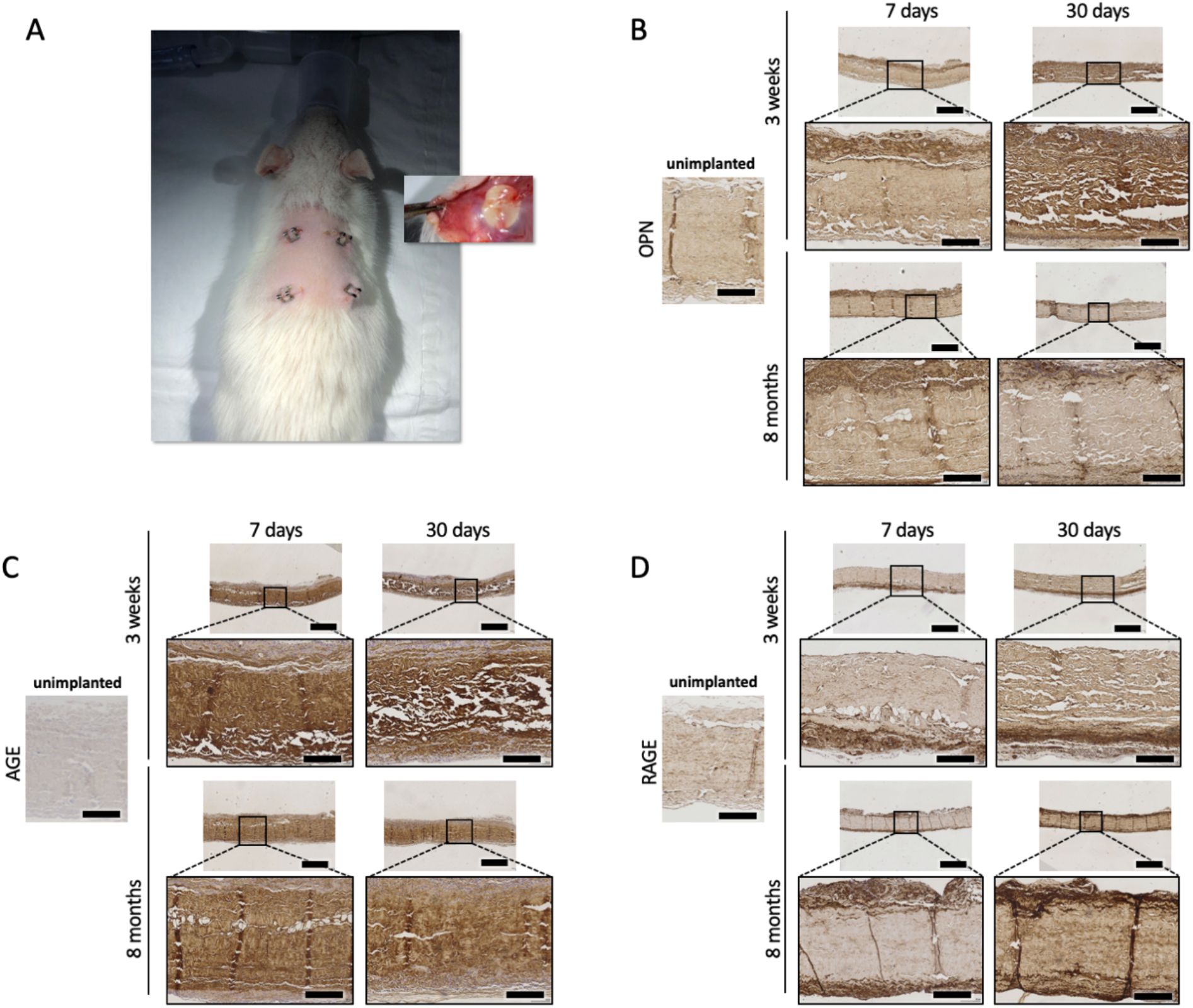
The rat subcutaneous implant model demonstrates accumulation of both tissue and circulating markers of mineralization and glycation. **A.** Photographs of the rat subcutaneous implant model immediately following BP implantation, and of BP during explantation. IHC for OPN (**B**), AGE (**C**), and RAGE (**D**) on BP following 7- or 30-day subcutaneous rat implantation in 3 week- and 8-month old rats (scale bars in lower magnification images represent 500 μm; scale bars in higher magnification images represent100μm).

**Supplemental Figure 3.**
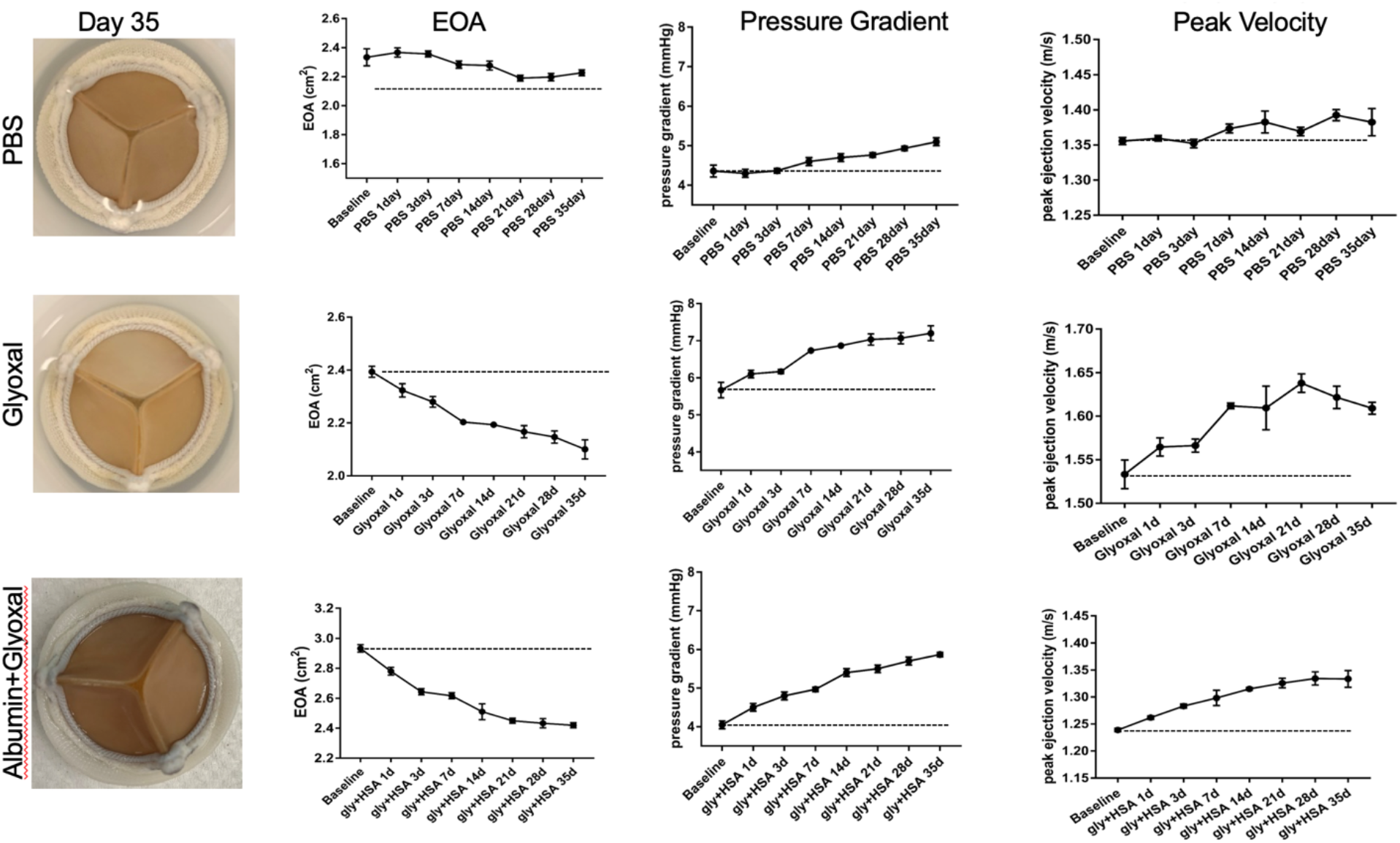
Analysis of hydrodynamic functions of BHV. Left: Visual images of three SAVR valves after 35-day *in vitro* incubation. Right: Values of parameters depicting hydrodynamic functions (EOA, mean pressure gradient, and peak ejection velocity) of the performances of three SAVR valves, measured by a pulse duplicator system, during the 35-day *in vitro* incubation period.

**Supplemental Figure 4.**
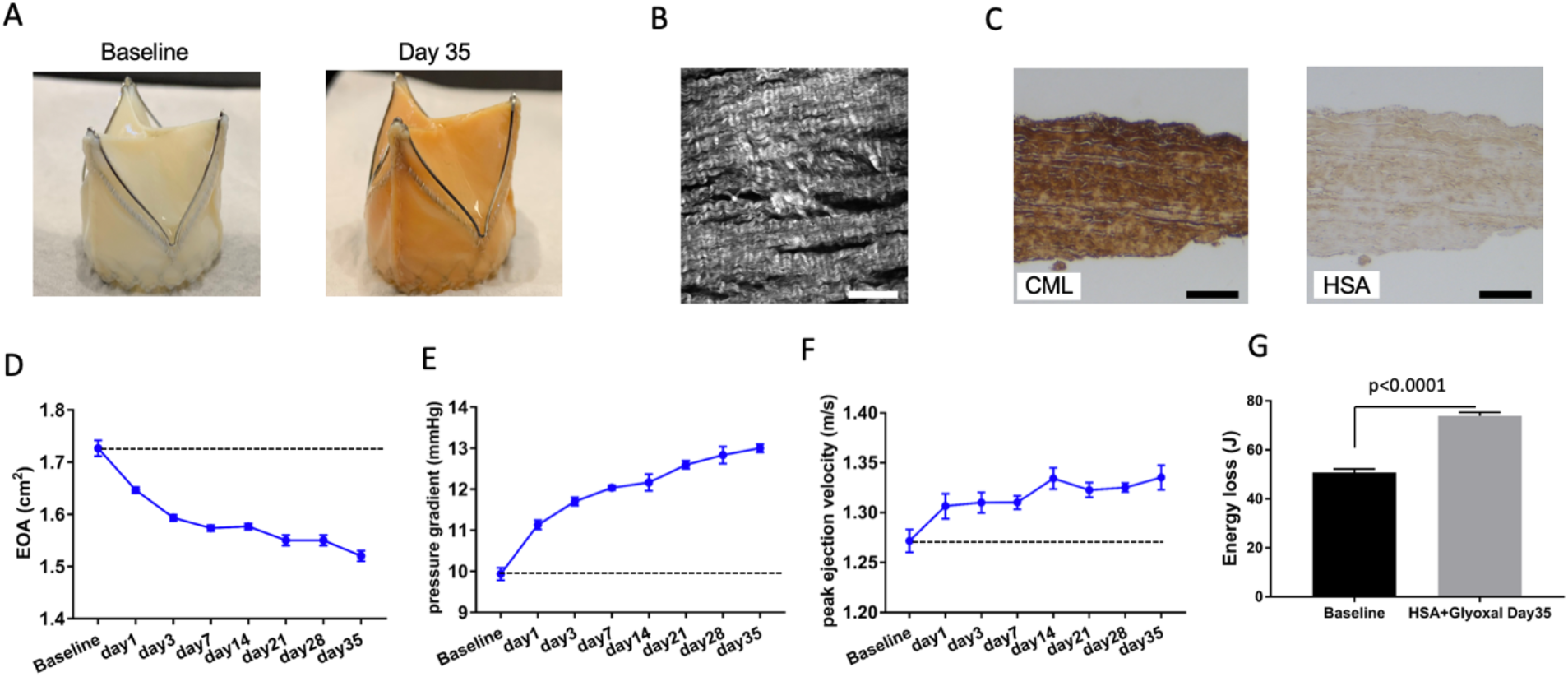
Impact of AGE and HSA accumulation on TAVR valve and its hydrodynamic functions. **A.** Visual images of an in-house unimplanted transcatheter bioprosthetic valve before and after 35-day *in vitro* incubation with HSA and glyoxal. **B.** SHG images of the TAVR valve after 35-day in vitro incubation (scale bar represents 100 μm). **C.** IHC staining for HSA, AGE, and CML on the TAVR valve after 35-day *in vitro* incubation (scale bars represent 250 μm). Values of parameters depicting hydrodynamic functions (EOA, **D.**, mean pressure gradient, **E.**, peak ejection velocity, **F.**, and energy loss, **G.**) of the TAVR valve measured by a pulse duplicator system during the 35-day *in vitro* incubation period.

